# Missing data in amortized simulation-based neural posterior estimation

**DOI:** 10.1101/2023.01.09.523219

**Authors:** Zijian Wang, Jan Hasenauer, Yannik Schälte

**Affiliations:** University of Bonn, Life and Medical Sciences Institute, 53115 Bonn, Germany; Helmholtz Center Munich, Computational Health Center, 85764 Neuherberg, Germany; Technical University Munich, Center for Mathematics, 85748 Garching, Germany

## Abstract

Amortized simulation-based neural posterior estimation provides a novel machine learning based approach for solving parameter estimation problems. It has been shown to be computationally efficient and able to handle complex models and data sets. Yet, the available approach cannot handle the in experimental studies ubiquitous case of missing data, and might provide incorrect posterior estimates. In this work, we discuss various ways of encoding missing data and integrate them into the training and inference process. We implement the approaches in the BayesFlow methodology, an amortized estimation framework based on invertible neural networks, and evaluate their performance on multiple test problems. We find that an approach in which the data vector is augmented with binary indicators of presence or absence of values performs the most robustly. Accordingly, we demonstrate that amortized simulation-based inference approaches are applicable even with missing data, and we provide a guideline for their handling, which is relevant for a broad spectrum of applications.

## 1 Introduction

Mechanistic models are used to describe and understand dynamical systems in a variety of research fields ranging from life and physical sciences to economics (N. A. Gershenfeld and N. Gershenfeld 1999; Kitano 2002). Commonly, these models depend on unknown parameters, which can be estimated by assessing the likelihood of observed data given parameters (Tarantola 2005; Raue et al. 2013). Classical parameter estimation methods (e.g. optimization, Markov-chain Monte-Carlo, approximate Bayesian computation (Raue et al. 2013; Pritchard et al. 1999)) are case-based. That is, they work on the level of individual data sets, such that the entire computationally expensive inference procedure needs to be repeated for every new data set.

However, often the same structural model is fitted to different data sets with potentially different parameters, e.g. to describe experiments under different stimuli, epidemic dynamics in different communities, or treatment response for different patients. In such cases, *amortized* inference methods are of interest. These first learn a mapping from synthetic data sets to e.g. likelihood or posterior distribution, which can subsequently be cheaply queried for many observed data sets (Papamakarios and I. Murray 2016; Lueckmann et al. 2021). A successful method, which is particularly applicable for the study of time series models, is BayesFlow. BayesFlow uses conditional invertible neural networks (cINN) to learn, conditioned on data, a reversible transformation from parameters to a tractable latent space, and has been shown to be superior to alternative approaches capable of amortized simulation-based inference, as well as to case-based methods when facing multiple data sets (Radev et al. 2020).

A problem persistent in many research areas is that data are incomplete, i.e. parts of the entries are missing. Possible reasons exist plenty, including incomplete entry, data loss, device malfunction, or study participant non-response. Further, in clinical studies measurements are often not taken at the exact time intervals or for different time spans, across patients or participants. There exist various missingness mechanisms as well as strategies to deal with them, e.g. by deletion or imputation (Enders 2022). In case-based inference, at its simplest, a likelihood or cost function can be formulated based on only the available data. However, amortized inference typically requires inputs to be of consistent structure and size across samples, and needs to know what entries are available, in order to learn the underlying data-parameter relationships correctly. For the specific case of data sets of different sizes, BayesFlow already allows to use summary networks yielding a fixed-size representation. However, this mechanism permits only e.g. time series of different length, but not entries to be missing at random intermediate points (see also the Supplementary Information, Section 1, for an illustration of how the established approach fails in this situation).

In this work, we propose and discuss three approaches of encoding missing data via fill-in values and augmenting the data. We integrate these into the BayesFlow workflow, and evaluate and compare their performance on three test problems. We find that an approach in which the data matrix is augmented with binary indicators of presence or absence of values performs the most robustly. Further, we demonstrate how this approach performs advantageously also in the simpler case of time series of different length.

### 1.1 Related work

There exist various possible missing data mechanisms, (not) missing (completely) at random (MCAR, MAR, NMAR, see Cheema (2014)). Our presented approach is agnostic of the underlying missingness mechanism, as long as the network learned to recognize what data are missing or present. Strategies to deal with missing data can be broadly divided into methods discarding entries with missing values, and approaches replacing missing values with imputed values. Imputation strategies include e.g. mean, maximum-likelihood, and multiple imputation, and deep learning approaches using e.g. LSTM networks (Song et al. 2020) and autoencoders (Khadka and Shakya 2020), see Nazabal et al. (2020) and Joel, Doorsamy, and Paul (2022) for a review. The faithful reconstruction of missing data using (multiple or weighted) imputation approaches relies on the accuracy of the used imputation method (see also a discussion of the inadequacy of overly simple, e.g. linear, imputation methods in the Supplementary Information, Section 2). Moreover, when properly accounting for uncertainty, imputation does not change the effective parameter posterior distribution, and therefore holds no conceptual advantage over approaches discarding missing data (see the Supplementary Information, Section 3). Instead, our approach falls into the first above category, i.e. it discards missing values. To the best of our knowledge, missing data discarding approaches for amortized inference have so far not been systematically studied.

## 2 Methods

### 2.1 Background

#### 2.1.1 Amortized simulation-based neural posterior estimation

A mechanistic model induces a likelihood function *θ* → *π*(*x*|*θ*) of measuring data 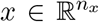 given model parameters 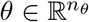. Parameter inference deals with the problem of estimating the unknown model parameters *θ*, given experimentally observed data 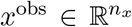. In a Bayesian setting, the likelihood is combined with prior information *π*(*θ*) on the parameters, giving by Bayes’ Theorem the posterior distribution

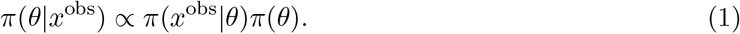

There are two major challenges to working with (1): (i) In many applications, the mechanistic model is only available as a simulator, allowing to generate synthetic data *x* ~ *π*(*x*|*θ*), but not to evaluate the likelihood function *π*(*x*^obs^|*θ*) (Tavaré et al. 1997; Jagiella et al. 2017). (ii) Often, the same model needs to be fitted to different data sets *x*^obs,*d*^, *d* = 1,…, *n_d_*, requiring the costly analysis of multiple posterior distributions.

BayesFlow approximates the posterior by a tractable distribution *π_ϕ_*(*θ*|*x*) ≈ *π*(*θ*|*x*) for any (*x*, *θ*) ~ *π*(*x*|*θ*)*π*(*θ*). The approximate posterior is parameterized in terms of a conditional invertible neural network (cINN) 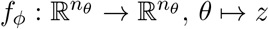, *θ* → *z*, conditioned on data *x*, which defines a normalizing flow (Papamakarios, Nalisnick, et al. 2021) between the posterior over the parameters *θ* and a standard multivariate normal latent variable *z*,

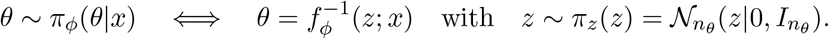

The neural network parameters *ϕ* are trained to minimize the Kullback-Leibler (KL) divergence between posteriors over all possible data sets *x*,

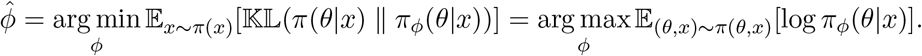

Via change of variable, it is *π_ϕ_*(*θ*|*x*) = *π_z_*(*f_ϕ_*(*θ*; *x*)) · |det *J_f_ϕ__*(*θ*; *x*)|, with straightforward calculation of the Jacobian 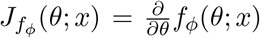 due to the cINN architecture. In practice, a Monte-Carlo approximation

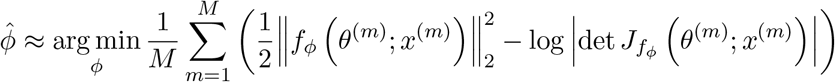

of the expectation is employed, with samples 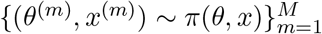.

#### 2.1.2 Summary networks

Instead of feeding the raw data directly into the cINN, summary statistics *h* = *h_ψ_*(*x*) can be employed. This has two advantages (Radev et al. 2020): First, tailored dimension reduction methods can adequately summarize redundant data and account for symmetries. Second, this enables the method to work with varying data set sizes 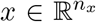 with random 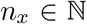, by transforming them into fixed-size representations. For example, an LSTM (Yu et al. 2019) can handle time series *x* of different length.

To avoid manual crafting, Radev et al. (2020) propose to learn maximally informative statistics from the data, by training the summary network parameters *ψ* jointly with the invertible network parameters, giving the joint objective

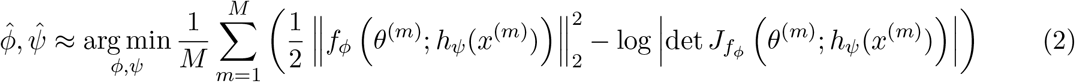

It can be shown that, provided sufficient training and expressiveness of *f_φ_* and *h_ψ_*, 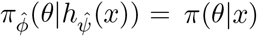 almost surely, i.e. the learned posterior perfectly approximates the actual one.

The upfront *training phase* can be expensive, as it might require many model simulations. Once the approximate posterior 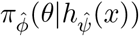 has been trained, for the *inference phase*, observed data *x*^obs^ are passed through the summary network, *h*^obs^ = *h_ψ_*(*x*^obs^). Then, latent variables 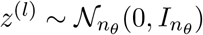, *l* = 1,…, *L* are sampled and transformed to samples 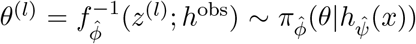 from the target distribution, by passing them through the cINN in inverse direction.

BayesFlow tackles both challenges (i) and (ii) above: It does not require likelihood evaluations and can thus be applied to any simulator. Further, it gives posterior approximations for any possible parameters and data. The inference phase is relatively cheap, as it does not require simulations of the mechanistic model, allowing to amortize the training phase when applied to different data sets *x*^obs^.

### 2.2 Encoding missing data

In many applications, not all datasets are complete. A prime example are clinical data, in which patients might have missed an appointment or dropped out of a study. In the case of incomplete data, the vector *x*^obs^ is not available completely, but some of its entries are missing, giving observed data 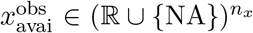. Explicitly, defining the binary availability mask *τ* ∈ {0, 1}^*n_x_*^, where 0 indicates absence and 1 presence of a data point, we consider 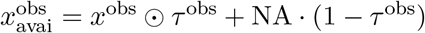, where ⊙ denotes the Hadamard product. There can be many causes and patterns of missingness, e.g. completely at random or dependent on the parameters. We are agnostic of the exact underlying mechanisms, and interested in the posterior 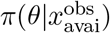 conditioned only on the available data. A different problem would be posed by the posterior *π*(*θ*|*x*^obs^) with *x*^obs^ the full data, which would however require a (faithful, multiple) reconstruction.

Missing values (“not available”, “not a number”) cannot be handled by neural networks, as they result in failing cost function and gradient evaluations. Simply dropping them from the data vector is no solution, as the context of which data were present would be lost. Further, the BayesFlow cINN *f_φ_* requires inputs of fixed size across samples. A summary network *h_ψ_* that permits inputs of variable size and transforms them into fixed-size representations constitutes one solution. This renders the method applicable to e.g. time series of different length, which can be mapped by an LSTM to a fixed-size latent state. However, this does not extend to random intermediate missing entries.

Here, we propose three ways of handling data sets *x*_avai_ with missing entries, to enable inference on the posterior *π*(*θ*|*x*_avai_). For simplicity, assume that the mechanistic model produces data sets of fixed size *n_x_*. This is in practice no restriction, as we can embed data sets of different size into one of maximum dimension 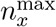, considering entries 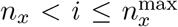 as missing too. We propose the following ways of encoding missing data into a vector *x*_aug_ such that the neural network can learn to detect and ignore them (see Figure 1 for an illustration of the resulting full workflow and examples):

- E1 (“Insert *c*”): Insert a constant value 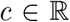 in place of missing data points. That is, set *x*_aug_:= *ι_c_*(*x*_avai_):= *x*_avai_ ⊙ *τ* + *c* · (1 – τ), i.e. *ι*(NA) = *c*, and *ι*(*x*) = *x* for 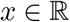.
- E2 (“Augment by 0/1”): As in E1, insert a constant value 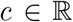 in place of missing data points. In addition, augment the data *x*_avai_ by the availability mask *τ* ∈ {0, 1}^*n_x_*^ to a combined matrix xaug:= [*ι_c_*(*x*_avai_), *τ*].
- E3 (“Time labels”): Restrict the data *x*_avai_ to available entries, yielding a reduced vector 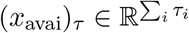. Augment this vector by a mask of time point indices 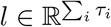, or alternative global identifiers, giving a combined data matrix *x*_aug_:= [(*x*_avai_)_τ_, *l*].

**Figure 1:**
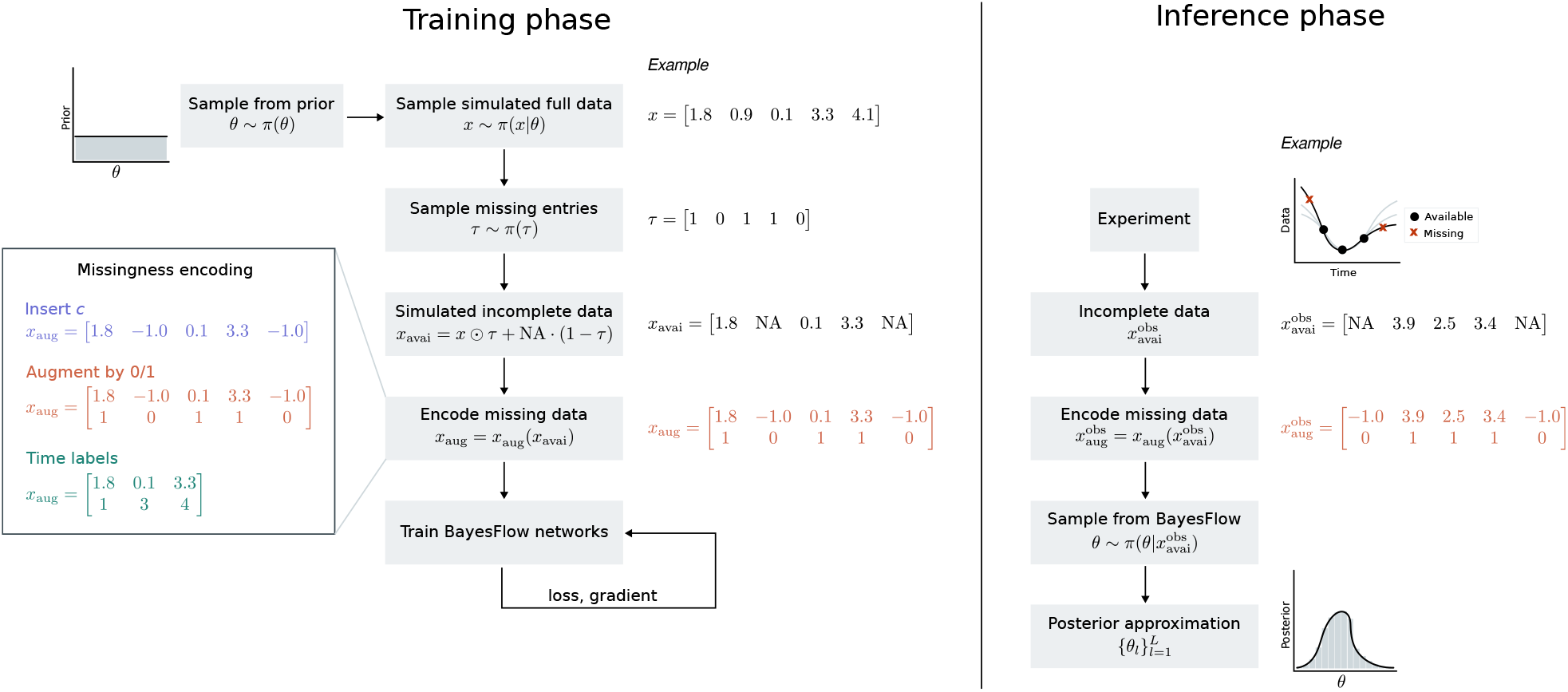
Illustration of the workflow combining BayesFlow with missing data encoding. Upfront training phase (left): Parameters *θ* ~ *π*(*θ*) are sampled from the prior to simulate complete data sets *x*_1:*N*_. Then, missing entries are randomly selected and encoded according to one of the three approaches “Insert *c*” (here *c* = −1), “Augment by 0/1” (here *c* = −1), and “Time labels”. The BayesFlow network is trained on such data sets with missing values using an online learning algorithm. Amortized inference (right): Experimentally observed incomplete data 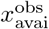 are processed using the preferred encoding approach (here “Augment by 0/1”, *c* = −1) and passed through the pre-trained BayesFlow network in its inverse direction. This leads to representative samples from the posterior conditioned on the available data 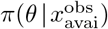. The upfront training amortizes over inference on arbitrarily many incomplete data sets.

### 2.3 Training algorithm with missing data

In order to facilitate recognizing the encoding of incomplete experimentally observed data during the inference phase, we train BayesFlow on data sets containing missing entries. Therefore, given complete simulated data *x*^(*m*)^ ~ *π*(*x*|*θ*), we explicitly simulate an availability pattern *τ*^(*m*)^ ~ *π*(*τ*) ∈ {0, 1}^*n_x_*^ in order to generate artificially incomplete data 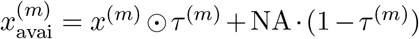. These are then passed through one of the missingness encoders, giving augmented data 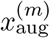, which are fed into summary and invertible network. The distribution of available entries *π*(*τ*) should incorporate any prior knowledge on missingness patterns, in order to train the network on realistic scenarios. At its simplest, we consider uniformly missing entries, i.e. with 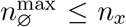 the maximum number of missing entries, we sample 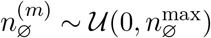, and then sample without replacement 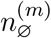 indices out of [1,…, *n_x_*].

In order to vectorize the propagation of samples through summary and invertible network, as well as gradient calculation via backpropagation, all samples within a batch must be of the same size. Thus, for approach E3 above, a single number *n*_∅_ of missing data points needs to be sampled per batch, like in the original BayesFlow implementation when considering time series of different length. The exact distribution of the *n*_∅_ missing entries over the data set can still be individual-specific. Meanwhile, in E1+2 the augmented data dimension is fixed and independent of the number of missing entries, such that the number of missing data points can be sampled for each sample individually. Sampling the number of missing data points on individual instead of batch level can be hoped to improve the stability of gradient descent when training the network parameters. This is because when using batch-level missingness sampling, the expected cost function varies over batches, that is we effectively use a different cost function in each step.

The entire algorithm for training and inference using BayesFlow with missing data is presented in Algorithm 1.

#### Algorithm 1 Amortized Bayesian inference for incomplete data using BayesFlow

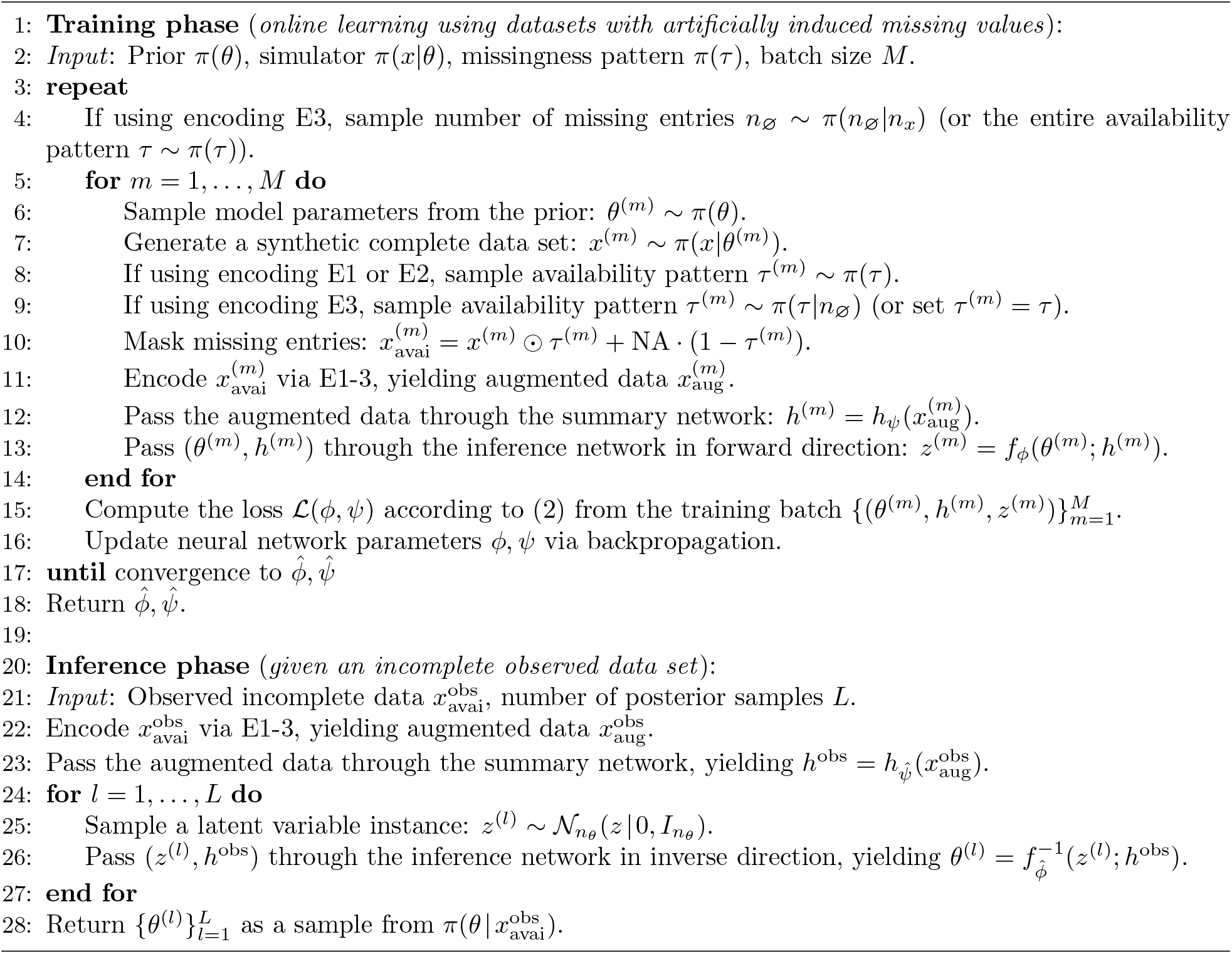

## 3 Results

We evaluated and compared the performance of the proposed missing data encodings E1-3 on three test problems – a simple conversion reaction model, an oscillatory model, and an SIR epidemiological model. Details on all test problems can be found in the Supplementary Information, Section 4. In the Supplementary Information, Section 5, we provide further analyses e.g. on convergence for all models, beyond the main results shown here in the main manuscript. When comparing results to the “true posterior”, we imply the distribution 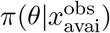 given only actually observed data, ignoring missing entries.

### 3.1 Implementation

Unless otherwise stated, we used the default settings of BayesFlow (version 0.0.0b1). For all problems, we ran the training phase for 300 epochs, each consisting of 1000 iterations, with one iteration denoting one batch of samples, over which the loss was calculated and backpropagation performed. The batch size was in most cases 128, except 64 for the SIR model. As summary network, we used an LSTM with the number of hidden units being a power of two close to the data dimension. The analyses were performed on a single CPU (AMD EPYC 7443 2.85 GHz) with 48 cores and 1 TB RAM. The full code underlying this study can be found at https://github.com/emune-dev/Data-missingness-paper, a snapshot of code and data is available on Zenodo at https://doi.org/10.5281/zenodo.7515458.

### 3.2 Conversion reaction model

We considered an ordinary differential equation (ODE) model of a simple conversion reaction *A* ⇌ *B*, a common building block in many biochemical systems and a widely used test problem (see e.g. Maier, Loos, and J. Hasenauer (2017) and Schälte and Jan Hasenauer (2020)). We assumed additive normally distributed measurement noise and that up to 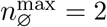 of the overall *n_x_* = 3 observations are missing. The aim is to infer the posterior of the two log-scale rate parameters *k*_1_, *k*_2_.

#### 3.2.1 All approaches perform well on simple test problem

For each of the encodings, we trained a 5-layer cINN with an LSTM with 8 hidden units as summary network. For E1+2, we used a fill-in value of *c* = −1, as the model only allows positive trajectories. For this simple problem, all three encodings led to accurate posterior approximations, even though the true posterior can be clearly non-Gaussian (Figure 2, Data set 1). In addition, in the special case that both values at *t*_2_ and *t*_3_ are missing and only the initial time point *t*_1_ is available, we are dealing with a completely uninformative data set, and all three approaches correctly returned the Gaussian prior distribution (Figure 2, Data set 2). This indicates that BayesFlow can conceptually comprehend each of the encodings, and focus on the available data points while ignoring masked missing entries.

**Figure 2:**
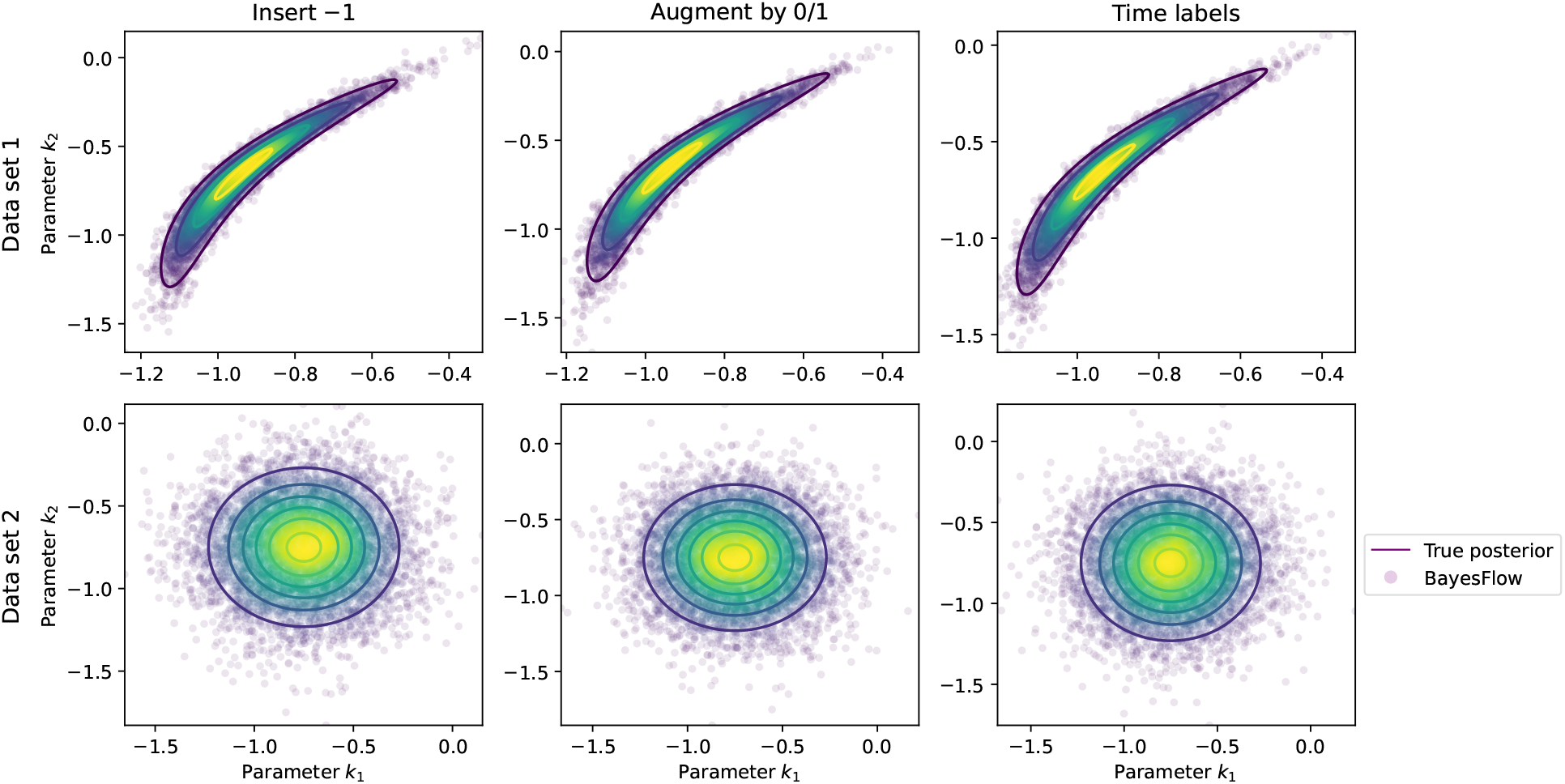
Posterior approximations for the Conversion reaction problem for data sets with missing entries. All three proposed encodings yield near-perfect approximations of the desired posterior given test data sets from this simple test problem. In Data set 2, no informative data are available, such that the posterior must equal the prior.

#### 3.2.2 Binary indicator augmentation increases robustness

In case no suitable dummy imputation value is known a-priori, e.g. when the model is flexible enough to simulate unbounded trajectories, when the prior range encompasses experimentally implausible regimes, or when noise levels can be large, the approaches “Augment by 0/1” and “Insert *c*” may have difficulties in distinguishing between signal and dummy imputation values representing missing data. Thus, we tested them on the conversion reaction model with an ambiguous dummy value of *c* = 0.5. We observed that the approach “Insert 0.5” misinterprets measured values in a neighborhood of 0.5 as missing data, whereas the approach “Augment 0/1”, additionally employing binary indicator augmentation, is capable of deciding correctly whether a value around 0.5 represents signal or missing value (Figure 3).

**Figure 3:**
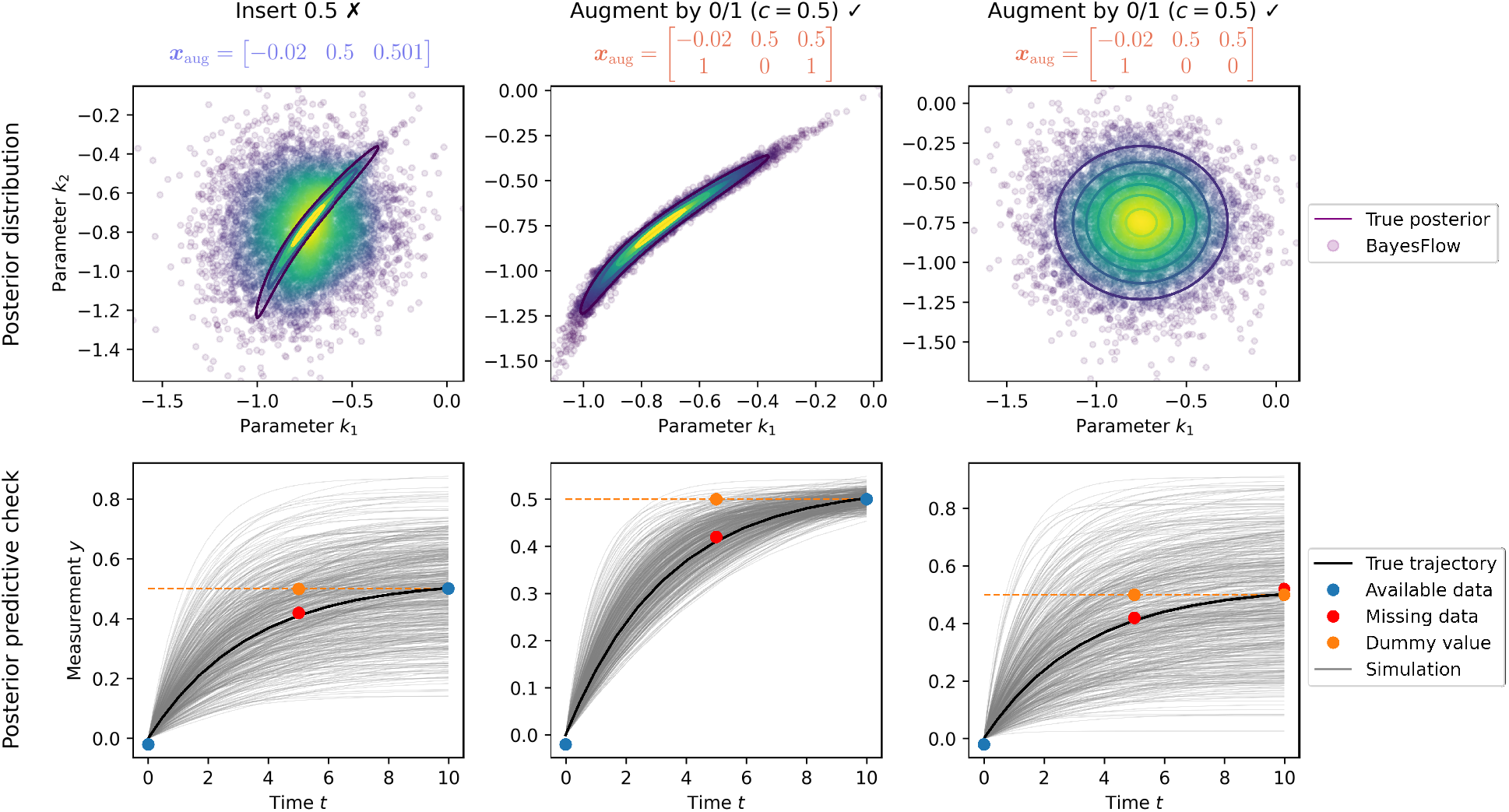
*Increased robustness through binary indicator augmentation in case of ambiguous dummy values (here: c* = 0.5). Left: The approach “Insert 0.5” sees a data set in which only the second observation is missing. However, the network misinterprets the signal 0.501 as another missing value. Hence, the estimated posterior is wrong, and the third available data point is not fitted by the re-simulated trajectories. Middle: The approach “Augment by 0/1” is able to correctly identify the value 0.5 in the second observation as a missing value and in the third observation as a signal. Consequently, the estimated posterior is correct, and the re-simulated trajectories fit the third data point, but not the second one. Right: The approach “Augment by 0/1” correctly interprets the value 0.5 in the second and third observation as missing value, although 0.5 would be a plausible data value for the third observation.

### 3.3 Oscillatory model

As a more complex example, we considered an oscillatory model, here a simple sinusoidal function sin(2*πa* · *t*) + *b*, in which we aim to infer frequency *a* and offset *b*. This is a prototype for oscillations frequently exhibited in biochemical systems, e.g. in the context of metabolism (Olsen et al. 2003) and cell cycle (Ingolia and A. W. Murray 2004). From a mathematical perspective, dynamic models producing oscillatory data are in general hard to fit, as the landscape of the cost function can be highly irregular and have multiple local minima (Pitt and Banga 2019).

#### 3.3.1 Improved performance on variable data set size as a special case of missing data

The original BayesFlow implementation (Radev et al. 2020) can already deal with time series models producing data sets of different length, by preprocessing the data with a suitable LSTM summary network, which reads in a given time series sequentially and summarizes it into a fixed-size representation, before feeding it into the cINN. Our missing data encoding also naturally covers this type of missingness, as time series of different length can always be interpreted as data in which the last 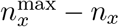 time steps are missing.

To study this, we assumed the number of observations in the oscillatory model to vary uniformly between 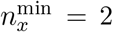 and 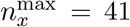. For both the original BayesFlow method and the approach “Augment by 0/1 (*c* = −5)”, we trained a 5-layer cINN jointly with an LSTM with 128 hidden units as summary network. Comparing the losses over 300 epochs (Figure 4a), we observed that for “Augment by 0/1” the loss converged faster, more smoothly and sometimes towards a lower final value, which corresponds to a better approximation of posterior distributions.

**Figure 4:**
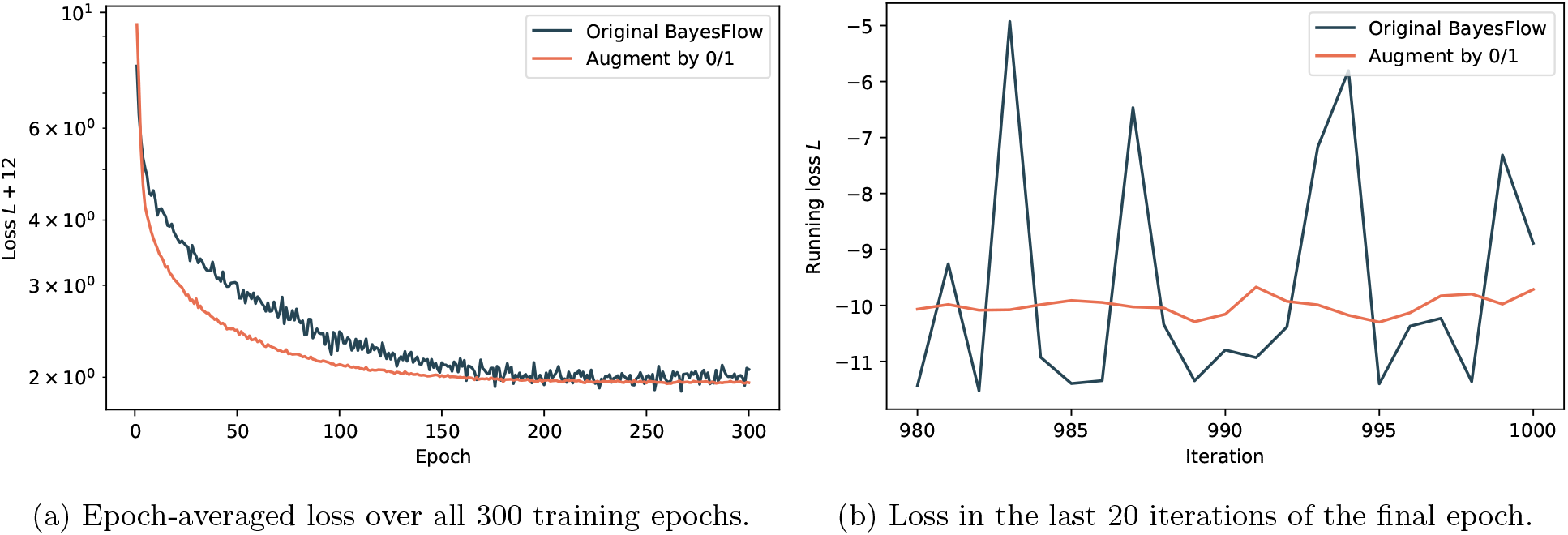
Comparison of loss behavior: In the case of different data set length, our missing data handling approach based on binary indicator augmentation achieves superior convergence to the original BayesFlow method, both globally (a) and on the level of individual iterations (b).

This can be explained by the fact that in the original BayesFlow method the number of available data points is sampled only once per batch, to enable vectorized operations. This gives a mathe-matically inexact approximation of the loss function, as its true value is only approximated as an average over iterations, but the network parameters are updated after each iteration. This leads to a loss that fluctuates severely from one iteration to another (Figure 4b), which affects the conver-gence of the stochastic gradient descent algorithm (Figure 4a). Contrarily, the approach “Augment by 0/1” yields augmented data sets of fixed size, thus it is possible to sample the available data points, or here rather the time series length, on the level of individual data sets and thereby target the same distribution in each iteration.

#### 3.3.2 Resampling missingness for each data set improves convergence

Assuming uniform missingness with a maximum number of missing time steps of 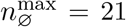, we compared the performance of the two encodings “Augment by 0/1” and “Time labels”. We trained a 5-layer cINN with an LSTM with 128 hidden units as summary network. A dummy value of *c* = −5 was employed for the former encoding.

Across test data sets generated for different ground truth parameters and exhibiting differently many missing values, we consistently observed the approach “Augment by 0/1” to approximate the true posterior better than the approach “Time labels” (Figure 5). The latter approach suffers, similar to the original BayesFlow algorithm in the previous section, from sampling the number of missing values only once per iteration and not per individual simulation. Consequently, its cost function exhibited substantially more fluctuations compared to “Augment by 0/1” (see the Supplementary Information, Figure S10). Similarly increased fluctuations could be observed when sampling the number of missing observations on batch level for the “Augment by 0/1” approach. However, in addition “Time labels” converged to an altogether worse cost function value. A similar cost function value was obtained by “Augment by 0/1” already after about 25 epochs. Comparing its posterior approximation at this early training stage with the final posterior approximation obtained by “Time labels”, we obtain qualitatively similar results (Supplementary Information, Figure S11). While further analyses would be needed, this could indicate that the network misinterpreted the time labels and could thus not converge to the true distribution.

**Figure 5:**
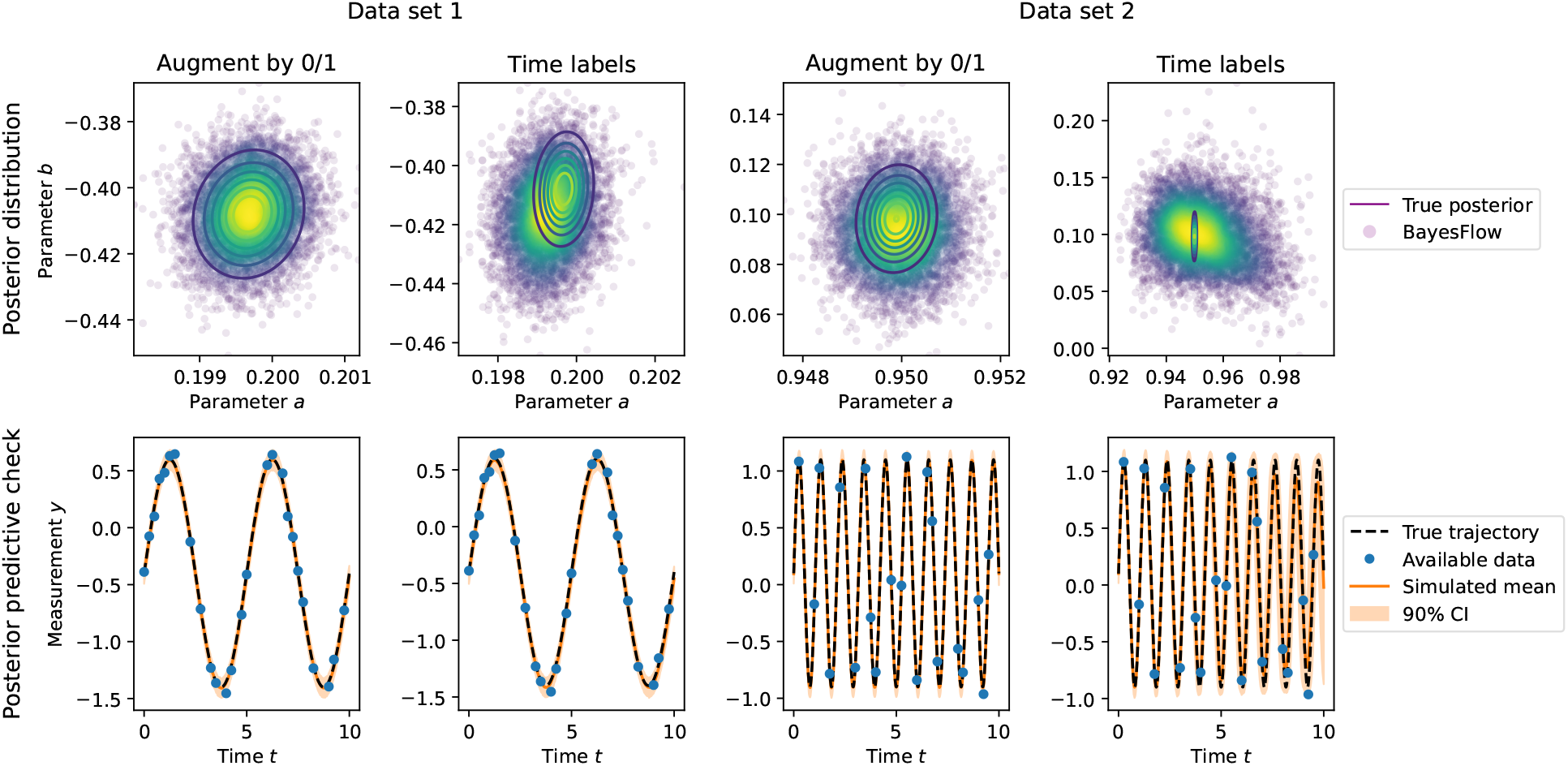
Results for the oscillatory model. Top: Posterior distributions. Bottom: Posterior predictive checks. Two data sets at parameters [0.2, −0.4] (Data set 1, left) and [0.95, 0.1] (Data set 2, right) are shown.

### 3.4 SIR epidemiological model

Compartmental models have been widely used to describe the course of the COVID-19 pandemic (Bertozzi et al. 2020; Raimúndez et al. 2021). However, infectious disease data are almost always subject to missing values. Therefore, we next studied a classic SIR model (Kuhl 2021), aiming to infer the log-scale transmission rate *b* and recovery rate *c*.

We assumed uniform missingness of at most 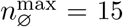 of the overall *n_x_* = 21 time steps, and trained a 5-layer cINN with an LSTM with 128 hidden units as summary network.

For the in previous studies most robust encoding “Augment by 0/1” with a dummy value of *c* = −1, we obtained precise approximations of the true posteriors, showing that our approach can deal with both more complex models and a high degree of missingness (Figure 6). See the Supplementary Information, Section 5.4 for “Insert −1”, which performed comparably, and “Time labels”, which converged slightly slower on this problem.

**Figure 6:**
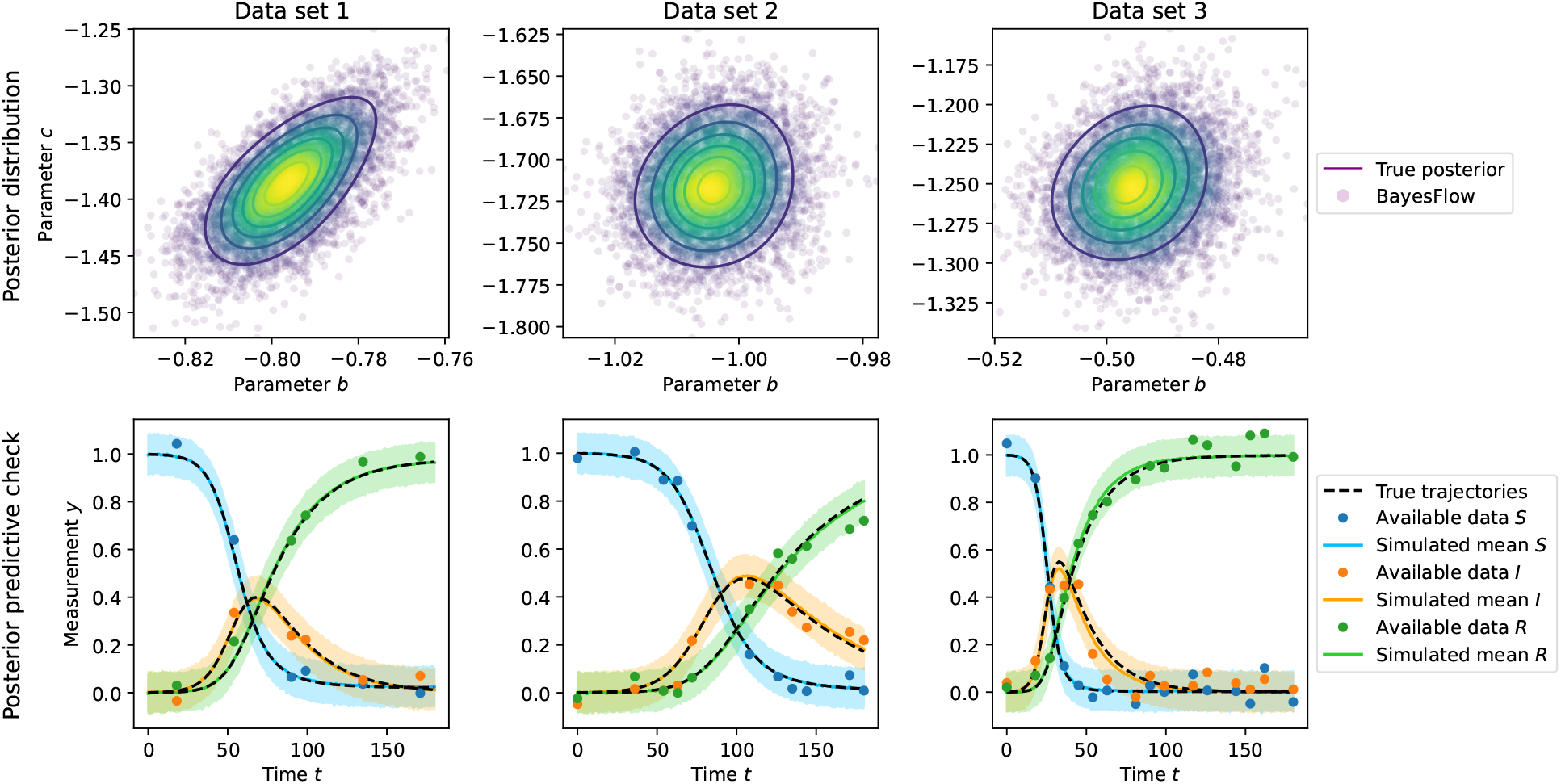
Results for the SIR model. Top: Posterior distributions. Bottom: Posterior predictive checks showing the means of noise-corrupted simulations and their centered 90% credible intervals. Three data sets at ground truth parameters [−0.8,-1.4] (Data set 1, left, *n*_∅_ = 15), [−1.0, −1.7] (Data set 2, middle, *n*_∅_ = 10) and [−0.5, −1.3] (Data set 3, right, *n*_∅_ = 5) are shown.

## 4 Discussion

Motivated by the fact that missing data are ubiquitous in experimental studies, in this work we presented approaches that allow to adequately handle them while performing inference over many observed data sets simultaneously. We achieved this by encoding the missing entries by fill-in values (“Insert *c*”) and augmenting the data by a binary mask indicating absence or presence (“Augment by 0/1”), or a mask identifying the available data points globally (“Time labels”), and using the established BayesFlow framework.

In particular, we found the approach “Augment by 0/1” to perform robustly across different problems. Unlike “Insert *c*”, it provides a binary indicator which can be easily interpreted by the neural network. This renders the approach particularly useful in case of ambiguous fill-in values. However, this comes at the cost of increased effective data set size. Thus, in cases a clear fill-in value can be found, “Insert *c*” may suffice and perform more efficiently. On the considered examples, we observed no substantial run time differences between the approaches.

In the original BayesFlow implementation, time series lengths were sampled only once per batch. This was in order to facilitate vectorized operations when propagating through the network. We showed that sampling the missingness pattern per individual, rather than once per batch, improves convergence, as it leads to a more stable Monte-Carlo approximation of the cost function. In particular, this renders “Augment by 0/1” a superior alternative to the “Time labels” approach, and also to the original BayesFlow implementation in the simplified case of time series of different lengths. The approaches are broadly applicable to various missingness scenarios. In particular, they allow also to handle irregular time series data, via projecting onto a high grid resolution with missing entries.

While we obtained some first promising results on how to combine amortized inference with missing data, several questions remain open: It remains to be studied how the suggested missingness encodings perform comparatively on more challenging application problems. Such problems would allow to realistically benchmark the approaches, e.g. on the trade-off of augmented data set size and accuracy. Instead of sampling the missingness pattern individually per model simulation, one could generate multiple data sets with missing entries from a single full simulation. Especially for expensive mechanistic models, this could be useful. However, the resulting trade-off of simulation cost and accuracy or convergence needs to be investigated. Further, while we here provided an integration into the BayesFlow methodology, the encodings may also prove useful for other amortized inference approaches, such as Tejero-Cantero et al. (2020), which however remains to be studied. Further, as we discussed briefly, an alternative to ignoring missing data is to impute faithful values. An investigation of the applicability of such approaches and a comparison to the here presented missingness encodings in terms of the obtained posterior approximation would be of interest.

In conclusion, in this work we presented and compared approaches to handle missing data in the amortized simulation-based neural posterior estimation framework BayesFlow. We believe that this will substantially improve its applicability on a wide range of problems.

## Acknowledgments

We thank Ullrich Köthe, Stefan Radev, and Marvin Schmitt for fruitful discussions, and Dilan Pathirana for assistance with the server setup.

