## Supplementary Information for "Missing data in amortized simulation-based neural posterior estimation"

### Supplementary information: Handling missing data in amortized likelihood-free neural posterior estimation

January 9, 2023

[1] University of Bonn, Life and Medical Sciences Institute, 53115 Bonn, Germany
[2] Helmholtz Center Munich, Computational Health Center, 85764 Neuherberg, Germany
[3] Technical University Munich, Center for Mathematics, 85748 Garching, Germany

### Contents

**1 Simple deletion of missing data does not work** **2**

**2 Linear imputation of missing data does not work** **3**

**3 Can imputation give more informative posteriors?** **4**

**4 Model details** **5**

### 1 Simple deletion of missing data does not work

The existing BayesFlow method allows to handle time series data of different length by converting them into a fixed-size vector of summary statistics using e.g. LSTM networks. In the following, we convince ourselves that the idea of simply deleting the missing values and passing the remaining set of available data points to the LSTM cannot work in the more general case when data may be missing at intermediate time steps.

To this end, we consider the conversion reaction model (see Section 4.1) with $N = 11$ observations from which we allow at most $N_{\varnothing}^{\max} = 6$ to be absent. We train a 5-layer cINN jointly with an LSTM with 32 hidden units on data generated according to the above stategy. Unsurprisingly, the so-trained BayesFlow network is not able to learn the correct posterior distribution, since the information on the specific position of missing observations is simply neglected. The misapproximated posterior for a test data set is illustrated in Figure S1.

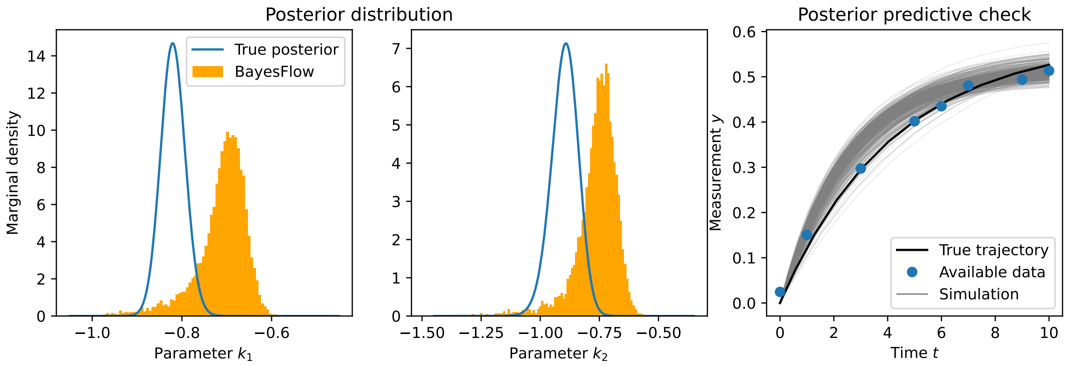

Figure S1: *Simple deletion of missing data cannot work.* For a test data set with missing values at $t = 2, 4, 8$, this overly simplistic approach leads to severe misapproximation of the true posterior distribution, which can also be seen from the re-simulated trajectories that fit the data poorly.

### 2 Linear imputation of missing data does not work

The goal of our proposed approaches is to estimate the posterior distribution $\pi(\theta|x^{\text{obs}}_{\text{avai}})$ conditioned only on the available data $x^{\text{obs}}_{\text{avai}}$. However, one might also be interested in the posterior $\pi(\theta|x^{\text{obs}})$ conditioned on the complete original data $x^{\text{obs}}$, since this distribution takes more information into account and is therefore more contracted. This type of inference would of course require a faithful imputation scheme $x^{\text{obs}}_{\text{avai}} \mapsto \tilde{x}^{\text{obs}}$ such that $\tilde{x}^{\text{obs}} \approx x^{\text{obs}}$.

We demonstrate that naive linear interpolation already fails for simple non-linear dynamics such as the conversion reaction model (see Section 4.1). Like in standard BayesFlow, we trained a 4-layer cINN on complete data sets. Then, for an incomplete test data set, we imputed the missing value from the available data via linear interpolation. The imputed data set was fed into the BayesFlow network trained on complete data, which resulted in misapproximated posteriors (Figure S2).

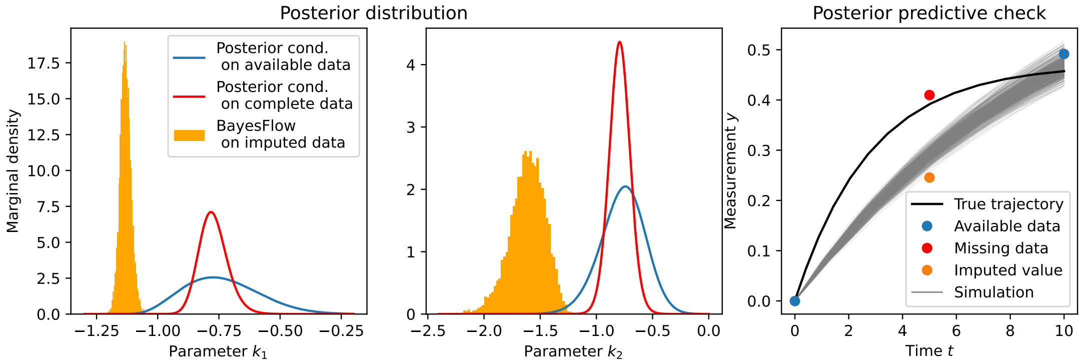

Figure S2: *Naive imputation of missing data based on linear interpolation cannot work.* Given is a test data set from the conversion reaction model with a value missing at $t = 5$. The posterior samples inferred from the imputed data by the BayesFlow network trained on complete data provide a poor approximation of the desired posterior conditioned on the complete data. The reason is that the linearly interpolated value is far off the true data value. Thus, the re-simulated trajectories are forced to fit the wrongly imputed value instead of the true value of the missing observation.

### 3 Can imputation give more informative posteriors?

The previous section underlines the importance of a reliable imputation scheme when attempting to recover the posterior $\pi(\theta|x^{\text{obs}})$ conditioned on the complete original data. In a Bayesian setting, however, we are less interested in a one-shot estimate of the posterior $\pi(\theta|\tilde{x}^{\text{obs}})$ given one specific set of imputed data $\tilde{x}^{\text{obs}} \approx x^{\text{obs}}$, which will in general be biased, but we should rather account for the uncertainty arising from the reconstruction of missing values $x^{\text{obs}}_{\text{avai}} \mapsto \tilde{x}^{\text{obs}}$.
Concretely, assume the imputation method returns a distribution $p(x^{\text{obs}}|x^{\text{obs}}_{\text{avai}})$ of complete data sets. For each such realization $x^{\text{obs}}$, the amortized posterior sampling method then gives a posterior distribution $\pi(\theta|x^{\text{obs}})$. To properly account for the uncertainty in the full data $x^{\text{obs}}$, we need to multiply the inferred parameter probabilities $\pi(\theta|x^{\text{obs}})$ with the according data probabilities $\pi(x^{\text{obs}}|x^{\text{obs}}_{\text{avai}})$ and marginalize over all possible full data values $x^{\text{obs}}$.
In this, either the imputation method is biased (if available, towards additional information), or it should be a faithful approximation $p(x^{\text{obs}}|x^{\text{obs}}_{\text{avai}}) \approx \pi(x^{\text{obs}}|x^{\text{obs}}_{\text{avai}})$, where $\pi(x^{\text{obs}}|x^{\text{obs}}_{\text{avai}})\pi(x^{\text{obs}}) = \pi(x^{\text{obs}}_{\text{avai}}, x^{\text{obs}})$ is the joint distribution of missing and complete data. That is, the imputation method recapitulates the generation of data and data missingness.
However, if we then integrate out all possible realizations of complete data, we find under the assumption $\pi(\theta|x^{\text{obs}}) = \pi(\theta|x^{\text{obs}}, x^{\text{obs}}_{\text{avai}})$:

$$\begin{aligned}\int \pi(\theta|x^{\text{obs}})p(x^{\text{obs}}|x^{\text{obs}}_{\text{avai}})\,dx^{\text{obs}} &= \int \pi(\theta|x^{\text{obs}}, x^{\text{obs}}_{\text{avai}})\pi(x^{\text{obs}}|x^{\text{obs}}_{\text{avai}})\,dx^{\text{obs}} \\ &= \int \pi(\theta, x^{\text{obs}}|x^{\text{obs}}_{\text{avai}})\,dx^{\text{obs}} \\ &= \pi(\theta|x^{\text{obs}}_{\text{avai}})\end{aligned}$$

This means we simply recover the posterior conditioned on the available data, whose estimation this paper is concerned with.
If instead $\pi(\theta|x^{\text{obs}}) \neq \pi(\theta|x^{\text{obs}}, x^{\text{obs}}_{\text{avai}})$, i.e. the missingness pattern contains information about the parameters beyond the complete data, we would still need to pass the missingness pattern to the amortized posterior sampling method, complicating its training. In doing so, we would however similarly finally obtain a marginalized posterior approximation

$$\int \pi(\theta|x^{\text{obs}}, x^{\text{obs}}_{\text{avai}})p(x^{\text{obs}}|x^{\text{obs}}_{\text{avai}})\,dx^{\text{obs}}) = \pi(\theta|x^{\text{obs}}_{\text{avai}}),$$

i.e. recover the posterior given the available data (including their missingness pattern) again.

### 4 Model details

In this section, we introduce the mathematical models that are used to test and compare the proposed methods for handling missing data.

#### 4.1 Conversion reaction model

We consider the conversion process

$$X_1 \underset{c_2}{\overset{c_1}{\rightleftharpoons}} X_2$$

with rate parameters $c_1, c_2 > 0$. If we denote the concentrations of the involved chemical species by $x_1$ and $x_2$, then their dynamics can be described by the following reaction rate equations:

$$\begin{pmatrix} \dot{x}_1 \\ \dot{x}_2 \end{pmatrix} = \begin{pmatrix} -c_1 x_1 + c_2 x_2 \\ c_1 x_1 - c_2 x_2 \end{pmatrix}$$

By specifying the initial value $(x_1(0), x_2(0)) = (1, 0)$, this linear system of ordinary differential equations has the unique analytic solution:

$$\begin{pmatrix} x_1(t) \\ x_2(t) \end{pmatrix} = \frac{1}{c_1 + c_2} \left[ \begin{pmatrix} c_2 \\ c_1 \end{pmatrix} + \begin{pmatrix} c_1 \\ -c_1 \end{pmatrix} e^{-(c_1+c_2)t} \right] \quad \text{for } t \geq 0$$

We assume that only the second state is measured, up to additive normal noise with known standard deviation $\sigma = 0.015$, i.e. the observation at time $t$ is given by:

$$y_t = x_2(t) + \varepsilon_t \text{ with } \varepsilon_t \sim \mathcal{N}(0,\, 0.015^2) \text{ independently in } t$$

The inference is performed for the log-scale parameters $k_j = \log_{10}(c_j)$ that are believed to follow normal priors:

$$k_1, k_2 \sim \mathcal{N}(-0.75,\, 0.25^2) \text{ i.i.d.}$$

The prior distribution is broad enough to allow sufficiently different dynamics, but also reasonably narrow so that it is unlikely to sample model parameters leading to very flat trajectories, in which case the measurement noise would be too dominant (Figure S3).
Depending on the application, we will assume, for the case of complete data, to observe data sets either consisting of $N = 3$ measurements at $t_0 = 0$, $t_1 = 5$ and $t_2 = 10$ or consisting of $N = 11$ measurements at $t = 0, 1, \ldots, 10$.

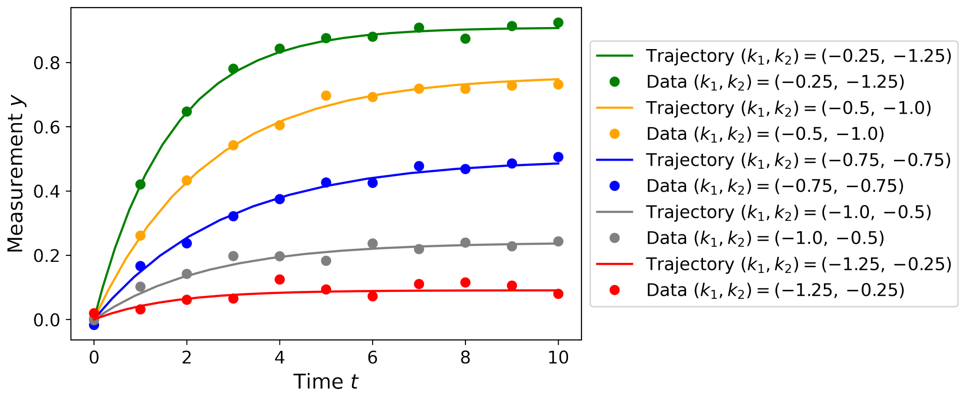

Figure S3: Conversion reaction model with $N = 11$ observations: Simulation of trajectories and noisy data for different parameters $(k_1, k_2)$ from the $2\sigma$-interval of their respective prior distribution

##### 4.2 Oscillatory model

Biochemical systems exhibiting oscillations are widely studied e.g. in the context of metabolism (Olsen et al. 2003) and cell cycle (Ingolia and Murray 2004). From a mathematical computational perspective, dynamic models producing oscillatory data are in general hard to fit, as the landscape of the cost function to be minimized can be highly irregular and have multiple local minima (Pitt and Banga 2019).

In our experiments, we consider the parametrized sine curve

$$x(t) = \sin(2\pi a t) + b$$

with frequency parameter $a$ and shift parameter $b$. The imposed prior distributions are as follows:

$$a \sim \mathcal{U}(0.1,\, 1), \qquad b \sim \mathcal{N}(0,\, 0.25^2)$$

We assume that at time $t$, the value of the sine curve can be observed up to some additive normal noise:

$$y_t = x(t) + \varepsilon_t \text{ with } \varepsilon_t \sim \mathcal{N}(0,\, 0.05^2) \text{ independently in } t$$

A complete data set should contain $N = 41$ observations at $t_k = \frac{1}{4}k$ for $k = 0, 1, \dots, 40$. Although seemingly harmless at first glance, this oscillatory model will already exhibit the important feature of a loss function that is difficult to optimize, so that convergence properties of different methods can be compared particularly well within the framework of this model.

##### 4.3 SIR epidemiological model

Compartmental models are nowadays a popular tool to describe and forecast the outcome of the COVID-19 pandemic (Bertozzi et al. 2020; Raimúndez et al. 2021). From a methodological viewpoint, such models are not only interesting because of their non-trivial dynamics, but also because their simulation requires at least solving an ODE.

For our purposes, it is sufficient to focus on the classic SIR model (Kuhl 2021). Let $S$ denote the number of susceptible, $I$ the number of infectious and $R$ the number of recovered individuals within a population of constant size $P = S + I + R$. Assuming the elementary reactions

$$S + I \xrightarrow{\beta} 2I, \qquad I \xrightarrow{\gamma} R \tag{1}$$

with transmission rate $\beta > 0$ and recovery rate $\gamma > 0$, the infection dynamics are quantified by the following non-linear system of ODEs:

$$\begin{aligned} \frac{\mathrm{d}S}{\mathrm{d}t} &= -\frac{\beta S I}{P}, \\ \frac{\mathrm{d}I}{\mathrm{d}t} &= \frac{\beta S I}{P} - \gamma I, \\ \frac{\mathrm{d}R}{\mathrm{d}t} &= \gamma I \end{aligned}$$

We specify the population size $P = 1000$ as well as the initial state $(S_0, I_0, R_0) = (999, 1, 0)$ and solve the initial value problem employing a standard ODE solver in Python with internal error control. The data at time $t$ are obtained by adding normally distributed measurement noise to the normalized state vector:

$$y_t = \frac{1}{P}\begin{pmatrix} S(t) \\ I(t) \\ R(t) \end{pmatrix} + \begin{pmatrix} \varepsilon_{t,1} \\ \varepsilon_{t,2} \\ \varepsilon_{t,3} \end{pmatrix} \quad \text{with } \varepsilon_{t,i} \sim \mathcal{N}(0,\, 0.05^2) \text{ independently in } t \text{ and } i$$

A complete data set contains $N = 21$ observations at $t_k = 9k$ for $k = 0, 1, \ldots, 20$. The inference is done for the log-scale parameters $b = \log_{10}(\beta)$ and $c = \log_{10}(\gamma)$ on which we impose normal priors:

$$b \sim \mathcal{N}(-1,\, 0.25^2),\; c \sim \mathcal{N}(-1.5,\, 0.25^2) \;\text{ independently}$$

### 5 Supplementary error analysis

In this section, we provide additional plots to illustrate the performance of our methods on the test problems.

#### 5.1 Methods

First, we briefly introduce the employed performance validation techniques.

- **Validation metrics**: The normalized root mean squared error NRMSE and the coefficient of determination $R^2$ are defined as follows for a sample of true parameters $\{\nu^{(m)}\}_{m=1}^{M}$ and a sample of estimated parameters $\{\widehat{\nu}^{(m)}\}_{m=1}^{M}$:

$$\text{NRMSE} := \sqrt{\sum_{m=1}^{M} \frac{\left(\nu^{(m)} - \widehat{\nu}^{(m)}\right)^2}{\nu_{\max} - \nu_{\min}}}, \qquad R^2 := 1 - \frac{\sum_{m=1}^{M}\left(\nu^{(m)} - \widehat{\nu}^{(m)}\right)^2}{\sum_{m=1}^{M}\left(\nu^{(m)} - \bar{\nu}\right)^2}$$

  Here, $\nu_{\max}$, $\nu_{\min}$ and $\bar{\nu}$ denote the maximum, minimum and mean of the true parameters, respectively. NRMSE measures how accurately the true parameter values are recovered by the estimates, and $R^2$ measures the proportion of variation in the sample of true parameters that is explained by the sample of estimated parameters. Perfect recovery is achieved when NRMSE $= 0$ and $R^2 = 1$.

  In our experiments, we choose $M = 500$, i.e. we compute the metrics on the basis of 500 test data sets. For the conversion reaction model, we compared the empirical means of the "true" posteriors obtained via MCMC sampling with the empirical means of the estimated posteriors from BayesFlow. For the oscillatory model, we compared the ground truth parameters (data generating parameters) with the empirical means of the estimated posteriors. This is justified as we consider a relatively high number of observations in the oscillatory model, even in the case of missing data, such that the ground truth parameters will give a good approximation of the true posterior means.

- **Simulation-based calibration**: Simulation-based calibration (SBC) exploits the insight that the Bayesian joint distribution is self-consistent (Talts et al. 2018). Concretely, it can be shown that averaging the exact posterior $\pi(\theta|x')$ over data $x'$ coming from the joint distribution $\pi(\theta', x') = \pi(\theta')\pi(x'|\theta')$ will recover the prior distribution (for a proof, see Appendix B in Radev et al. (2020)):

$$\pi(\theta) = \iint \pi(\theta|x')\pi(\theta', x')\, d\theta' dx'$$

  If the exact posterior is replaced by an inadequate approximation, the above equality will be violated. Such violations can be detected by computing rank statistics which compare a sample of data generating prior parameters with the according approximate posterior samples and inspecting the resulting histogram for uniformity (SBC plot). While uniform SBC plots indicate good approximation, different types of deviations from the uniformity allow interpretations such as over-/underfitting or a systematic bias in the approximate posteriors. For illustrative examples of such deviations and their interpretations, we refer to Talts et al. (2018).

  In our test problems, we computed the SBC histograms using 5000 prior parameters (5000 data sets) and 250 posterior samples per data set.

#### 5.2 Conversion reaction model

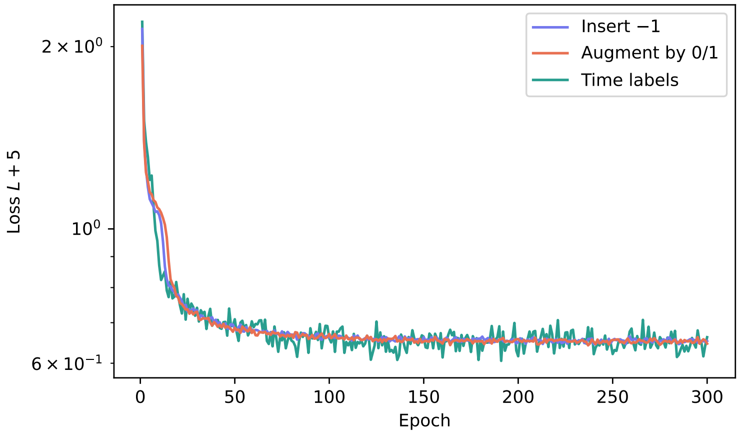

Figure S4: *Convergence plot for the conversion reaction model.* For this simple test model, all three proposed approaches result in a similar convergence rate of the loss function. The loss curve for the "Time labels" approach is more jumpy due to the sampling of missingness on the batch level.

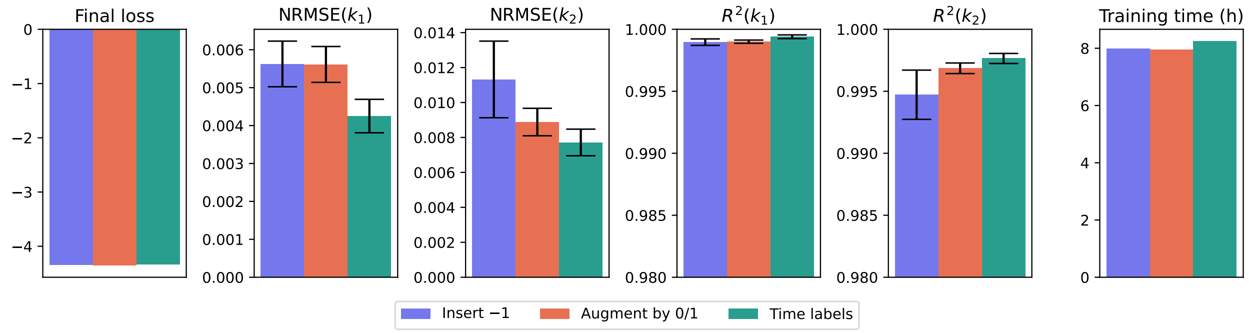

Figure S5: *Error metrics for the conversion reaction model.* The three encodings lead to similar error metrics in terms of the final loss as well as NRMSE and $R^2$ scores. The "Time labels" approach performed marginally superior on this problem. The training time is approximately the same for all three encodings.

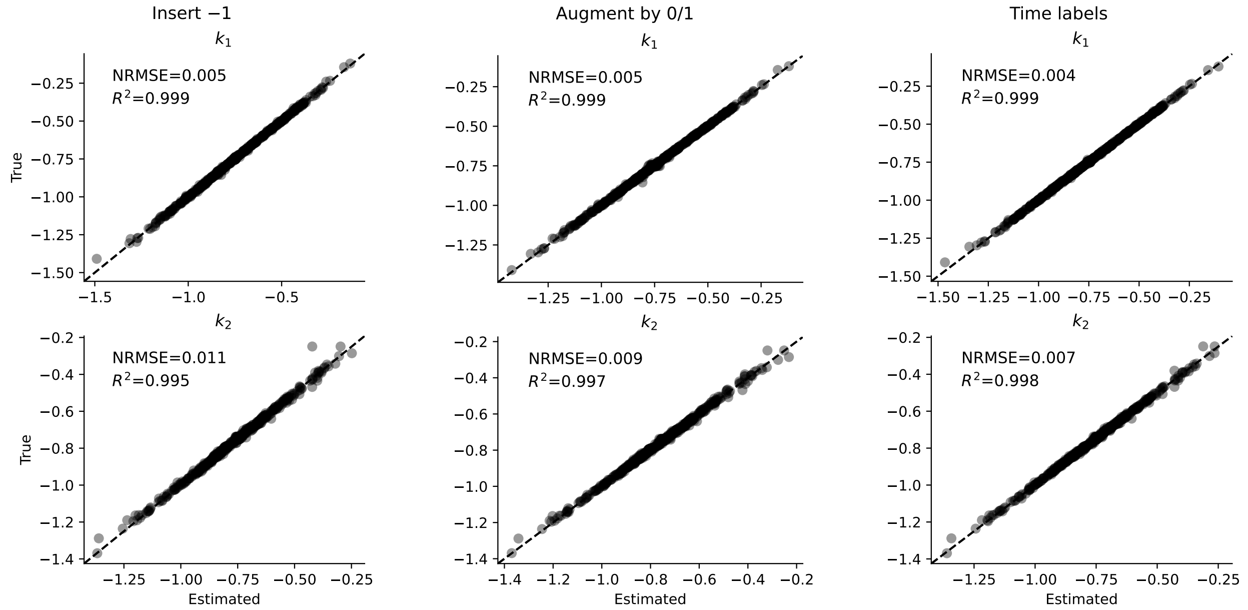

Figure S6: *True vs. estimated for the conversion reaction model.* Posterior means estimated by the BayesFlow network are in great accordance with the "true" means obtained via MCMC sampling, showing that all three encodings perform very well on this test problem. Marginal differences as in Figure S5 result from a tiny portion of data sets.

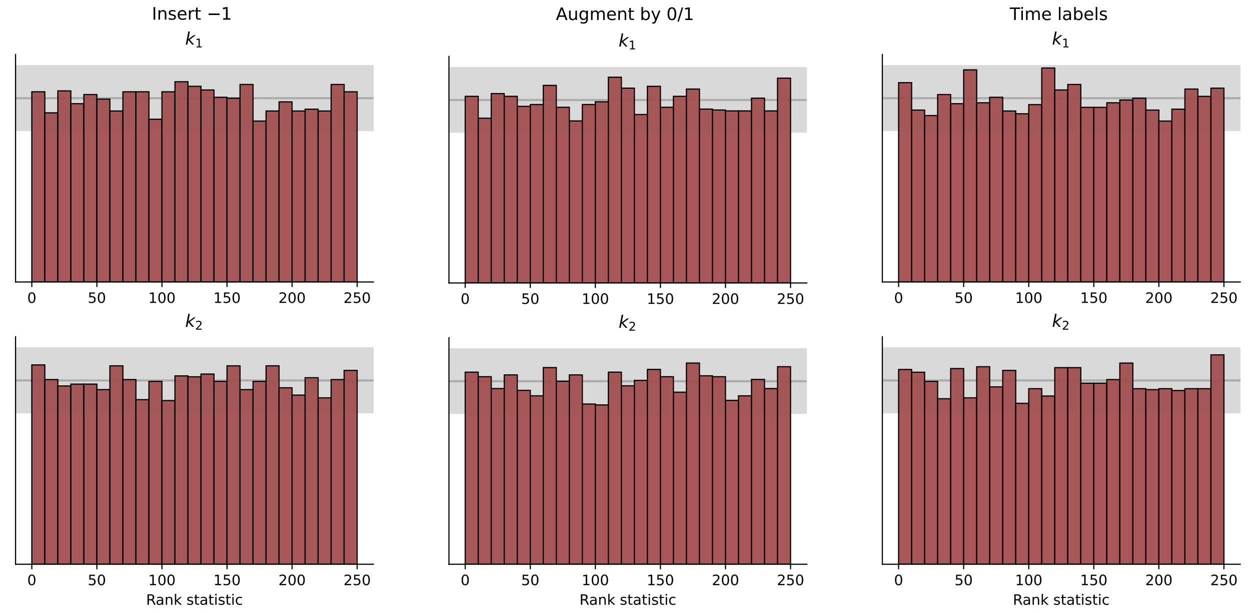

Figure S7: *SBC for the conversion reaction model.* The histograms exhibit uniformity, indicating the networks have converged and no systematic bias or over-/underfitting of the posteriors is detected.

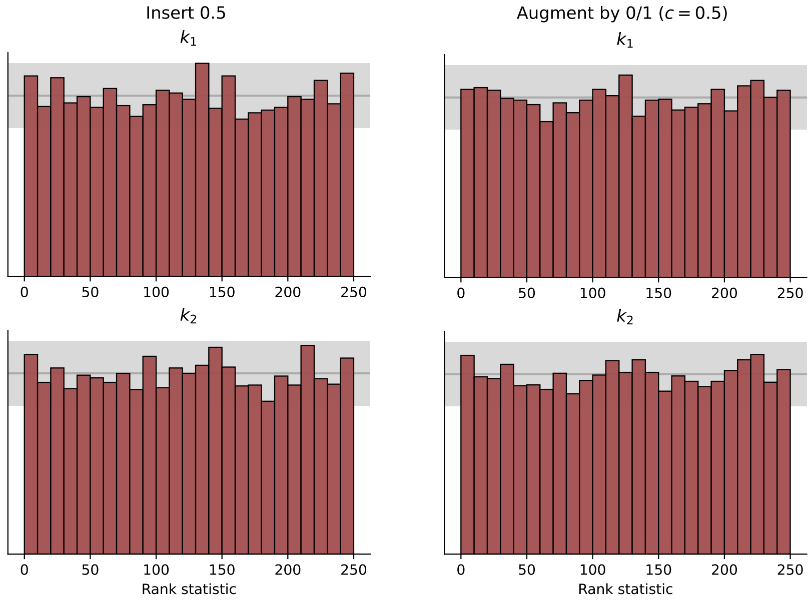

Figure S8: *SBC for the conversion reaction model (investigating ambiguous dummy imputation values).* No systematic bias or over-/underfitting of the posteriors is detected for the trained networks.

#### 5.3 Oscillatory model

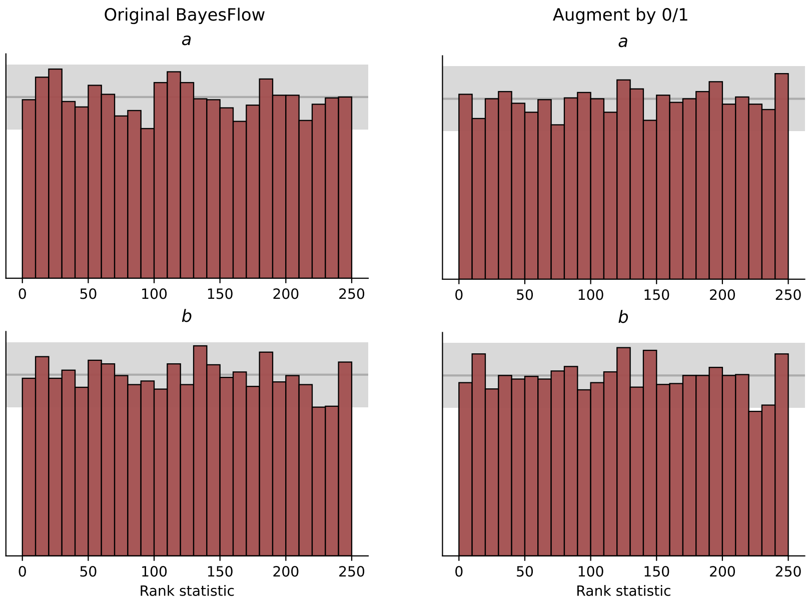

Figure S9: *SBC for the oscillatory model (variable data set length).* No systematic bias or over-/underfitting in the posteriors is captured by the histograms.

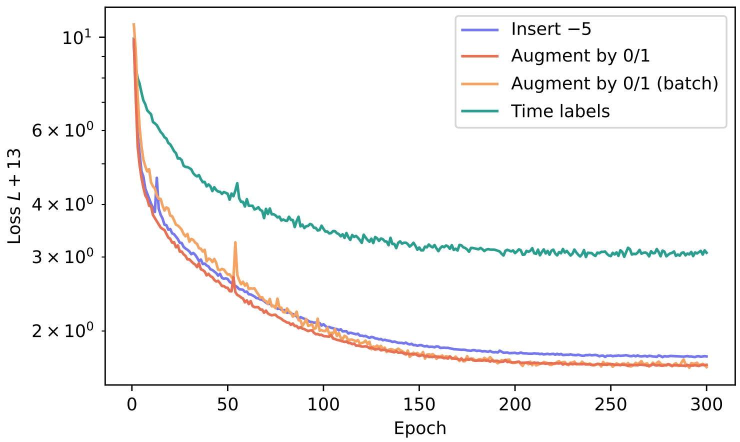

Figure S10: *Convergence plot for the oscillatory model.* The approach "Augment by 0/1" performs the most robustly: It achieves a lower final loss than "Insert $-5$", although $c = -5$ is an unambiguous dummy value for this model. The approach "Time labels" converges very poorly. This behavior cannot be merely explained by the sampling of number of missing observations on batch level, as it cannot be reproduced by using the binary augmentation with batch sampling ("Augment by 0/1 (batch)"). It seems that in the case of oscillatory data, the network may misinterpret the time labels and thus not converge properly.

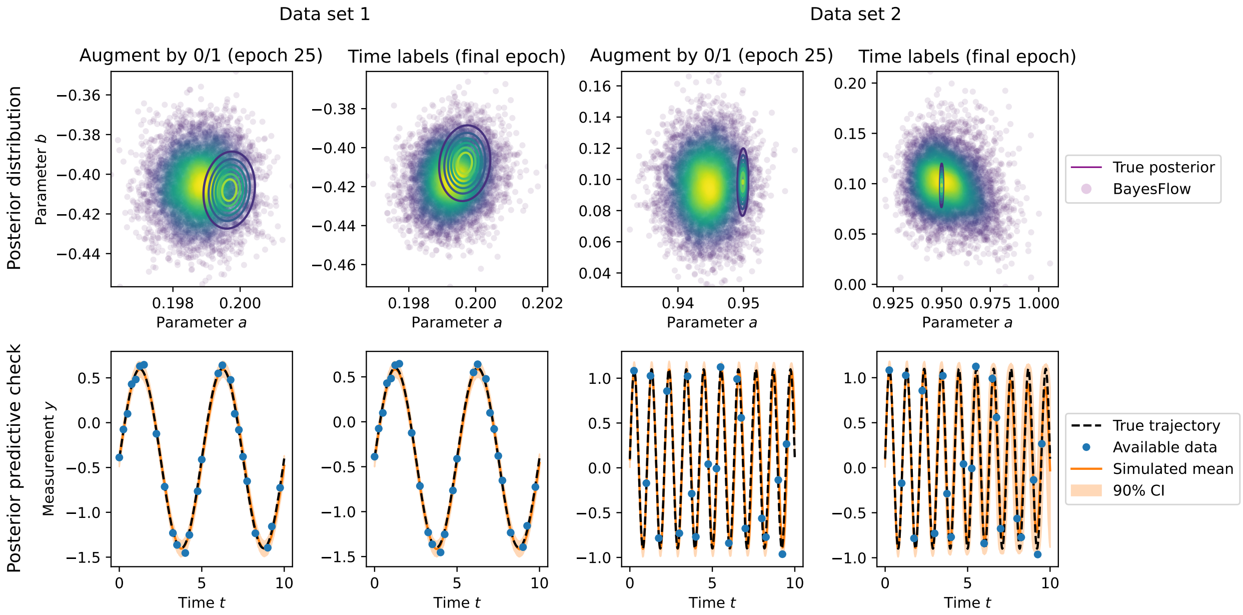

Figure S11: *Posterior estimation and predictive check for the oscillatory model, comparing "Time labels" and "Augment by 0/1" in an earlier generation.* The final loss achieved by the network trained with the encoding "Time labels" is approximately reached by the network trained with "Augment by 0/1" already in epoch 25. Here, we compare the posterior approximation of "Augment by 0/1" at this early stage of training with the final posterior approximation obtained by "Time labels". We observe similarly poor approximations of the posteriors.

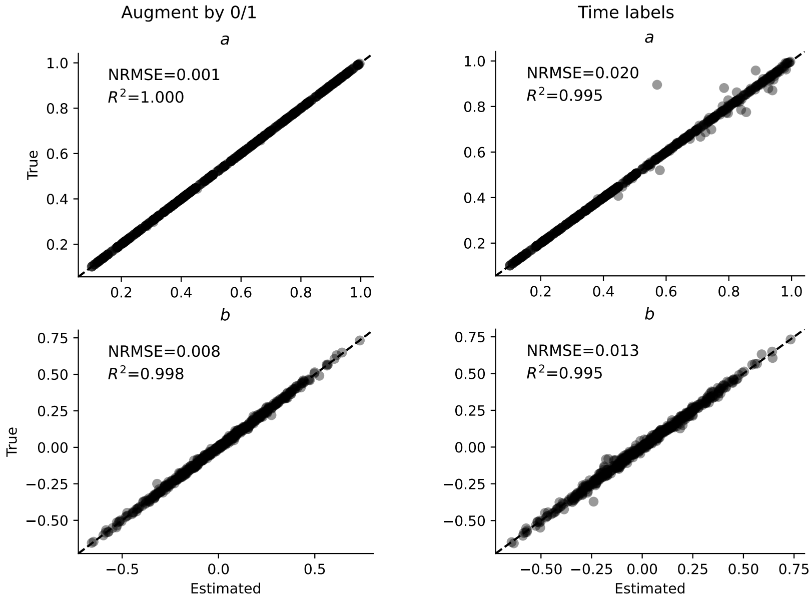

Figure S12: *True vs. estimated for the oscillatory model.* The encoding "Time labels" leads to larger deviations of the estimated means from the ground truth than the encoding "Augment by 0/1".

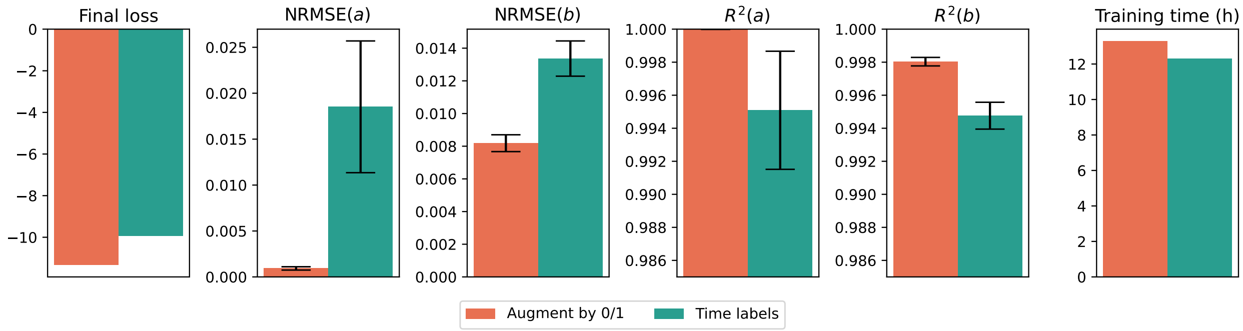

Figure S13: *Error metrics for the oscillatory model.* In accordance with Figure S12, the error metrics NRMSE and $R^2$ indicate a much better approximation of the ground truth parameters by the approach "Augment by 0/1" than by "Time labels".

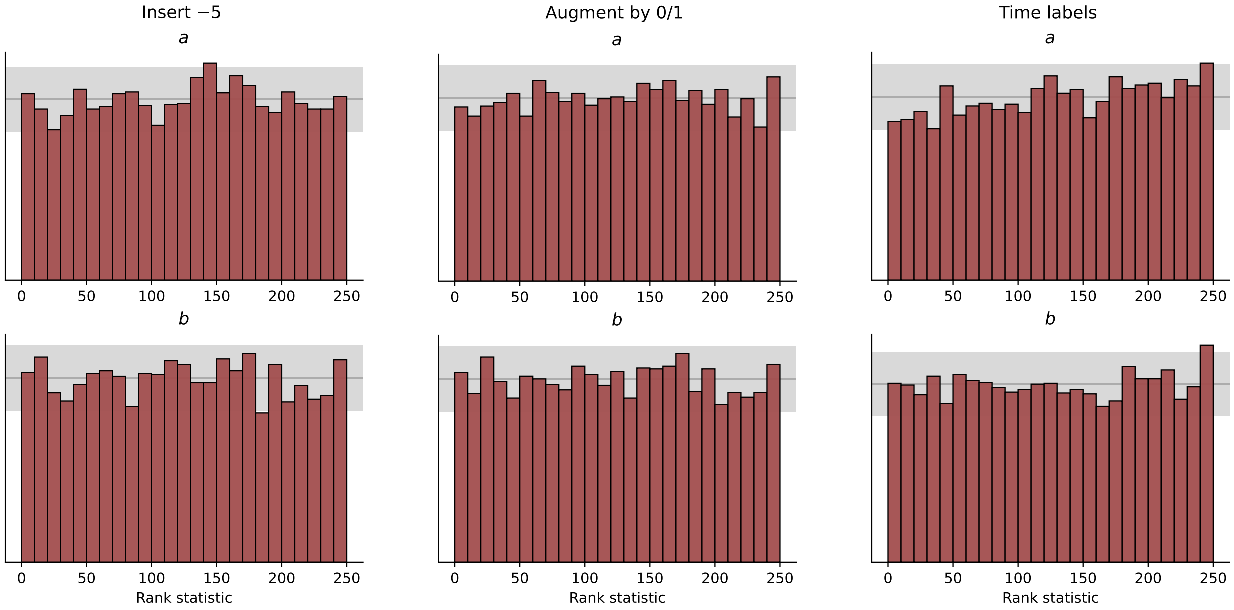

Figure S14: *SBC for the oscillatory model.* No clear systematic bias or over-/underfitting in the posteriors is detected by the histograms.

#### 5.4 SIR epidemiological model

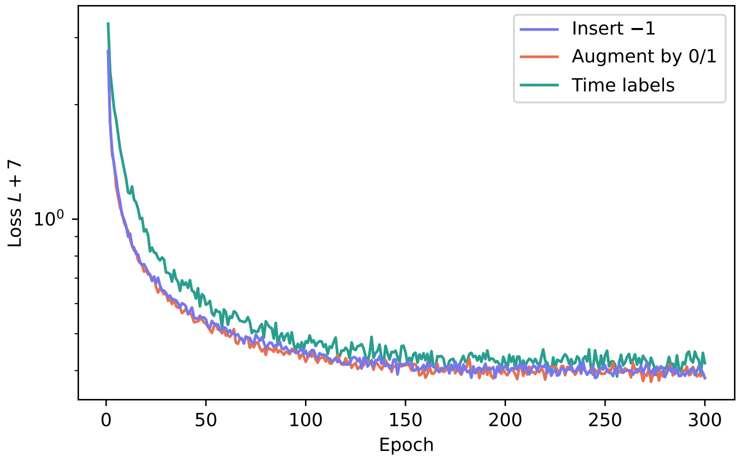

Figure S15: *Convergence plot for the SIR model.* The losses of the networks trained with the encodings “Augment by 0/1” and “Insert −1” behave similarly. They converge faster, more smoothly and towards a lower final value than the loss for “Time labels”.

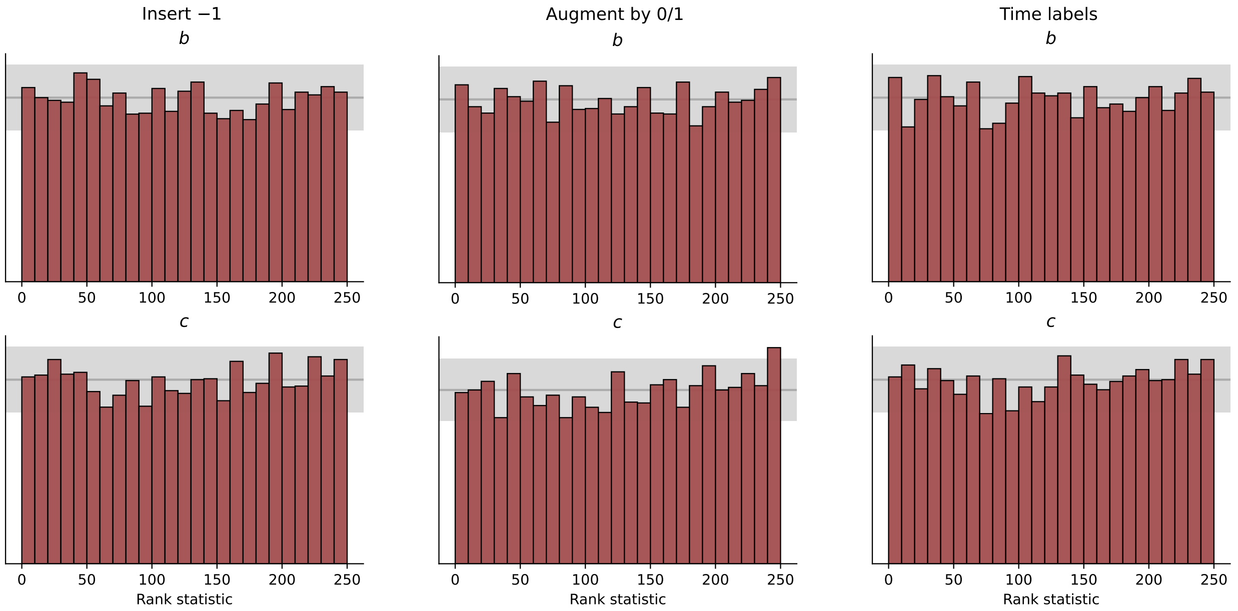

Figure S16: *SBC for the SIR model.* No clear systematic bias or over-/underfitting in the posteriors is detected by the histograms.

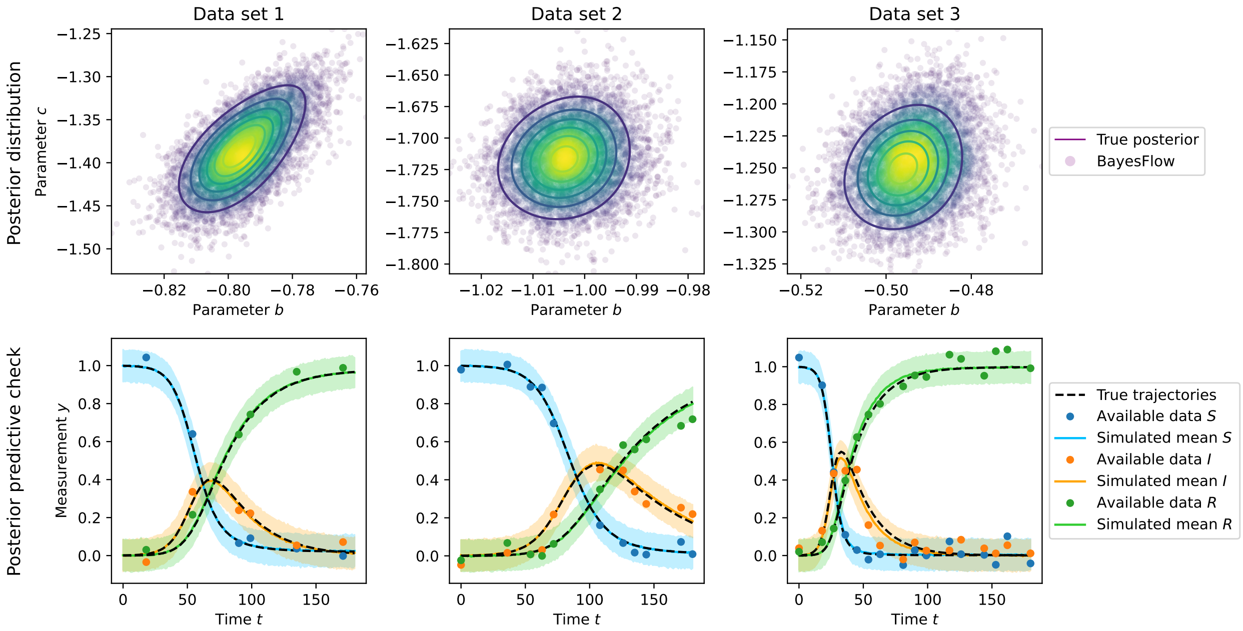

(a) Posterior samples and predictive checks using “Insert −1”

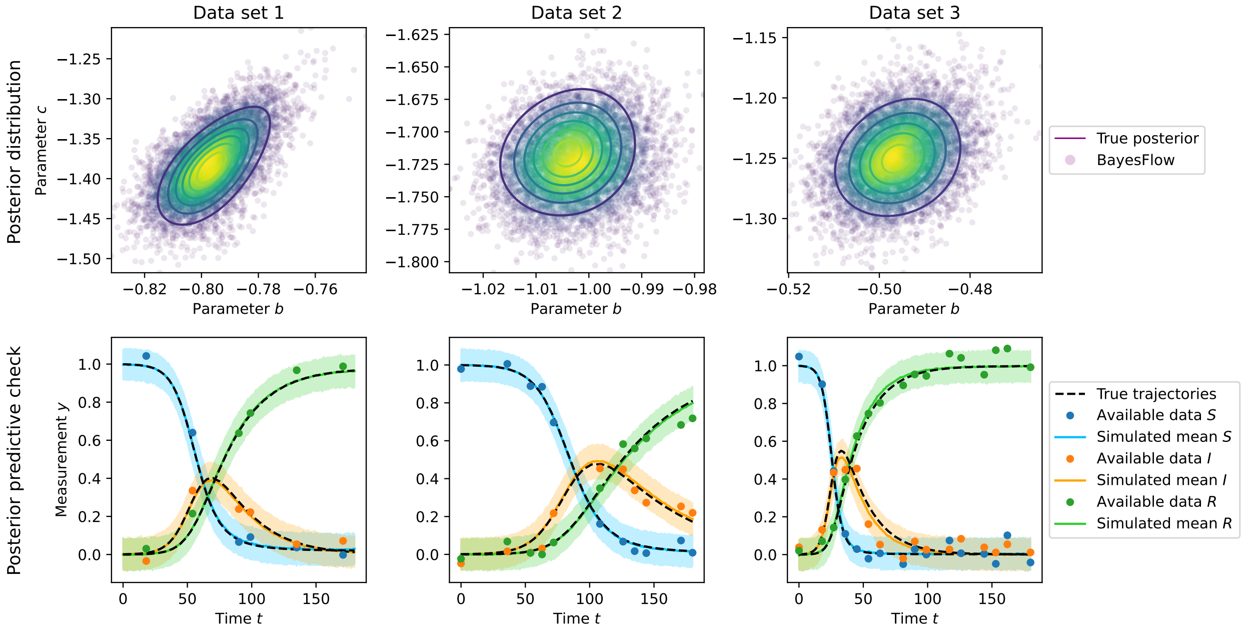

(b) Posterior samples and predictive checks using “Time labels”

Figure S17: *Results for the SIR model using the encodings (a) “Insert −1”, (b) “Time labels”.* Respective top: Posterior distributions. Respective bottom: Posterior predictive checks showing the means of noise-corrupted simulations and their centered 90% credible intervals. Three data sets at ground truth parameters $[-0.8, -1.4]$ (Data set 1, left, $n_\varnothing = 15$), $[-1.0, -1.7]$ (Data set 2, middle, $n_\varnothing = 10$) and $[-0.5, -1.3]$ (Data set 3, right, $n_\varnothing = 5$) are shown.
